# The macroecology of passerine nest types, in the light of macroevolution

**DOI:** 10.1101/360958

**Authors:** Jay P. McEntee, Zoe Zelazny, J. Gordon Burleigh

**Affiliations:** Department of Biology, University of Florida, Gainesville, FL 32611, USA

**Keywords:** Evolution, behavior, function, predation, thermoregulation, macroecology, trait macroevolution

## Abstract

Passerine birds build a diversity of nests to lay and incubate eggs, and to house nestlings. Open cup, dome, and hole (or cavity) nests have distinct advantages and/or disadvantages related to predation risk and thermoregulation. We used macroecological and macroevolutionary approaches to test contrasting predictions from considering these consequences. Patterns of prevalence across latitude and elevation for the roofed nest types (holes and domes) provide no evidence that their thermoregulation benefits promote colonization of colder environments. These patterns are more consistent with the role of predation in determining where dome-nesting species in particular occur. Macroevolutionary analyses suggest that diversity patterns for nest types along major ecological gradients mostly arise from how clades with conserved nest types have diversified across gradients, rather than arising from local adaptation. Lastly, we reveal a negative relationship between body mass and latitude in hole-nesting passerines, which runs counter to Bergmann’s rule.

**Statement of authorship:** JPM and JGB designed the study. JPM and ZZ compiled data from the literature. JPM performed statistical analyses with input from JGB. JPM and JGB wrote the manuscript, and ZZ contributed to revisions.

**Data accessibility statements:** Data were obtained from existing sources in the literature, cited in the manuscript.

## Introduction

Across animal diversity, many species construct nests, modifying their environments to carry out particular activities. The passerine birds, members of the order Passeriformes, are among the most familiar nest-builders. The great majority of passerine species build nests in which they lay and incubate eggs, and subsequently house altricial nestlings (Hansell 2000). Some passerines additionally use nests for roosting (Kendeigh 1961), although this behavior is far less widespread across diversity. The diversity of nest sites, construction materials, and architecture among passerines has made this group a preferred study system for the ecology and evolution of nest building (Collias & Collias 1984; Collias 1997; Hansell 2000; Price & Griffith 2017).

While there is great diversity in nest forms and sites among passerine species, a number of authors have categorized passerine nests into three basic types: hole/cavity nests (hereafter hole nests), dome nests, and open cup nests (Wallace 1868; Studer 1994; Martin 1995; Collias 1997; Martin *et al*. 2017). Both hole and dome nests are roofed nests, distinct from open cup nests, which are open above. Whereas hole nests are constructed within some existing roofed structure, dome nests are constructed in the open. The two primary functions of passerine nests are thought to be protection from predation and thermoregulation. Roofed nests are thought to be advantageous in both respects (Nice 1957; Collias & Collias 1984; Lamprecht & Schmolz 2004; Auer *et al*. 2007; Martin *et al*. 2017) The advantages of open cup nests may be that they are less time-consuming or less energetically expensive to construct (Mainwaring & Hartley 2013).

The consensus from the literature appears to be that hole nests have the lowest predation rates of the three nest types (Nice 1957; von Haartman 1957; Skutch 1966; Ricklefs 1969; Collias & Collias 1984; Oniki 1985; Martin & Li 1992; Martin 1993, 1995; Auer *et al*. 2007), with dome nests generally having lower predation rates than open cup nests (Oniki 1979; Linder & Bollinger 1995; Auer *et al*. 2007; Martin *et al*. 2017). The relatively longer developmental periods of eggs and nestlings in roofed-nesting species have been viewed as evidence of adaptive evolution to lower predation rates, with open cup nesting species forced to shorten development periods because of high predation rates (Martin 1993, 1995).

Evidence indicates that some roofed nests aid nest inhabitants by providing greater thermoregulatory benefits, with temperatures inside roofed nest buffered relative to external temperatures (Lamprecht & Schmolz 2004). Roofed nests might further provide protection from damaging insolation (Collias 1964) and precipitation (Collias & Collias 1984). These thermoregulatory benefits reduce the amount of energy consumed in thermoregulation (Kendeigh 1961; Buttemer *et al*. 1987), potentially benefitting roofed nesting species in many different environments. The roosting of passerines in hole (Kendeigh 1961) and domed nests (Skutch 1961; Buttemer *et al*. 1987) during cold nights outside the breeding season provides stark evidence for the thermoregulatory benefits of enclosed nests in cold environments. By comparison, non-breeding season roosting in nests is exceedingly rare among open cup-nesting species (Skutch 1961). The smaller body size of dome-nesting passerines, in contrast to both hole- and open cup-nesting passerines, has further been claimed as evidence for the thermoregulation benefits of domed nests in cold environments (Collias & Collias 1984; Martin *et al*. 2017), as smaller-bodied animals lose heat more rapidly (Calder 1983).

The relative advantages of roofed nests in terms of predation and thermoregulation yield predictions about where roofed-nesting species should be most prevalent along environmental gradients. Evidence generally indicates that nest predation rates are higher at tropical latitudes (Ricklefs 1969; Oniki 1985) and possibly in the southern temperates (Martin 1996; Martin *et al*. 2017), as compared to higher latitudes and the northern temperates. If nest predation rates help determine the geographic ranges of species, we predict that both hole- and dome-nesting species should be relatively more prevalent at lower latitudes. Thermoregulatory pressures could yield a number of different latitudinal patterns, but we focus on one prediction here: the ability of enclosed nests to slow heat loss for eggs and altricial, featherless nestlings suggests they should be especially helpful in cold environments where ambient temperatures are far below the temperatures necessary for egg development. Thus, we expect that enclosed nests should be relatively more prevalent at extremely high latitudes where the warmest seasons are still cold. Comparisons of community-level data have reported higher frequencies of dome-nesting at low latitudes and in the southern hemisphere (Auer *et al*. 2007; Martin *et al*. 2017), consistent with the expectations from nest predation rates, and not with thermoregulation pressures. Latitudinal trends in hole-nesting are less frequently reported in the literature, although some evidence suggests that passerine hole-nesting is less frequent in tropical forest than in the northern temperates (Ricklefs 1969), consistent with expectations from thermoregulation pressures. Barve and Mason (2015), however, found no correlation between cold breeding conditions and the evolution of cavity nesting in the Muscicapidae using phylogenetic logistic regression.

Existing evidence consistently shows that predation rates decrease with elevation in the tropics (Skutch 1985, Boyle 2008, Jankowski et al. 2013). Jankowski et al. (2013) hypothesized that relaxed predation pressures at higher tropical elevations might lead to the evolution of life history characteristics unsuitable for high predation pressures at lower elevations. If predation rates are consistently higher in open cup nests relative to hole and dome nests (Nice 1957; Snow 1978; Oniki 1985; Hall *et al*. 2015; Martin *et al*. 2017), nest types could evolve under these dynamics, leading to high frequencies of cup nests at high elevations. Alternatively, if the thermoregulatory demands of high-elevation environments are more important than predation in shaping the nest type use across elevations, we would predict that enclosed nests (dome and hole nests) should attain high frequencies at higher elevations. This should be especially true in the tropics, where temperature differences at different elevations are more consistent across annual cycles (Janzen 1967; Londoño *et al*. 2017) – i.e. there are no warm seasons at >3000m elevation that allow species to nest at temperatures similar to the balmy lowlands. Thus, elevation within the tropics should be a more consistent proxy for breeding temperatures than latitude. To our knowledge, little previous work has explored variation in passerine nest type frequency along elevational gradients. However, intraspecific variation in nest material, construction, and placement consistent with adaptation to cold temperatures in the Hawaiian honeycreeper *Hemignathus virens* have been found (Kern & Van Riper III 1984).

Nest type patterns along gradients could result primarily from adaptation to environmental conditions if nest type is labile, or primarily from the differential diversification of clades dominated by different nest types if nest type is conserved. To investigate which of these mechanisms is responsible for the nest type patterns we see, we must reconstruct the evolutionary history of these nest types across the passerines. While behavioral traits have often been considered to be especially labile (Darwin 1874; Blomberg *et al*. 2003), a recent analysis indicated that passerine nest type may not be (Price & Griffith 2017).

To further contextualize the evolution of passerine next types, we examined the association of nest types with body size evolution. Body size is thought to be associated with different nest predation rates, with larger birds suffering higher nest predation rates (Brightsmith 2005), and thermoregulatory pressures, where heat loss is a greater concern for smaller species (Calder 1983). Collias and Collias (1984) suggested that the small size of dome-nesting species provides support for the importance of thermoregulation and/or protection from abiotic environment in roofed nests, and that these thermoregulatory benefits could be especially important in cold environments at high latitudes. Dome-nesting species have been found to be consistently smaller than open cup-nesting species in community-level analyses across regions (Martin et al. 2017), evidence viewed as indirect support for the thermoregulation functions of domed nests given that heat loss increases with surface:volume ratios.

## Methods

### Nest type scoring and data set

We scored nest types for the 4,373 passerine species whose nest type or nesting behavior was adequately described to assign a score in the Handbook of the Birds of the World Alive (del Hoyo *et al*. 2015, last accessed 30 June 2016, hereafter HBW Alive). These 4,373 species represented 74.0% of the 5,912 passerine species in the HBW Alive taxonomy. We categorized 96.6% of these species’ nests as open cup, domed, or hole (we use the term “hole” to refer to any nest built inside a tree cavity, rock crevice, or earthen bank). In distinguishing between open cup and domed nests in ambiguous cases, as for nests described as ‘purses’, we scored nests as ‘open cup’ where descriptions or photographs indicated that nests are exposed above. In cases where nests described as “purses” have side entrances and are not open above, they were scored as “domed.” We scored nests described as “partially domed” as “domed.” The remaining 3.4% of the species were scored either as nesting in more than one nest type category or as brood parasites, which do not construct a nest or incubate eggs. We refer to the data set that includes all 4,373 species as the “all species” data set.

### Phylogeny

In order to account for the history of nest type evolution in our macroecological analyses, we reconstructed the history of nest type transitions across the passerine phylogeny. For this purpose, we used the topology of the supermatrix phylogenetic tree of Burleigh et al. (2015). We transformed the branch lengths to be ultrametric by performing a penalized likelihood analysis with r8s v. 1.71 (Sanderson 2003). The size of the phylogenetic tree rendered a more complex Bayesian approach, e.g. BEAST (Drummond & Rambaut 2007) computationally infeasible. The branch lengths were calibrated using twenty fossil calibrations from throughout the avian phylogeny (Baiser *et al*. 2017). The optimal smoothing parameter was estimated in r8s via a cross-validation analysis. For this analysis, the age of crown Psittacopasserae was fixed to 60 million years, midway between the minimum (53.5 my) and maximum (66.5 my) estimated ages. We determined the optimal smoothing parameter by checking how closely the unconstrained fossil age estimates matched their fossil-constrained age estimates, resulting in an optimal smoothing parameter of 3.2. We then trimmed the phylogenetic tree so that it included only the Passeriformes.

Ancestral state reconstruction required taxonomic reconciliation between the trait data set (del Hoyo *et al*. 2015, last accessed 30 June 2016) and phylogenetic tree (Clements *et al*. 2011; Burleigh *et al*. 2015). We identified potentially mismatched taxa using the name.check function from the R package geiger (Harmon *et al*. 2008). We examined all cases where a species in the Burleigh et al. (2015) phylogenetic tree did not have a corresponding species with the exact same name in the data set using the HBW Alive taxonomy. We examined the taxonomic history for these species in Avibase (https://avibase.bsc-eoc.org/avibase.jsp?lang=EN), and changed the species name to match the Burleigh et al. (2015) phylogenetic tree when an alternate species name matched a taxon name from the HBW Alive taxonomy. Many of these cases involved either the use of different genus names or alternate spellings. Taxa treated as subspecies in the HBW Alive (2015) taxonomy and species in Burleigh et al. (2015) were not included in our analyses.

There were 3,242 passerine species with nest decriptions that could be matched to the tips in the Burleigh et al. (2015) phylogenetic tree (hereafter the “parsimony” data set). These included species scored as using only one of the three nest type categories (hole, cup, and dome), as well as species nesting in more than one nest type (hole or cup, hole or dome, cup or dome), and brood parasites. We believe that estimating the transition rates among the seven nest types using maximum likelihood methods is unwise, as some transitions are too rare to justify rate estimation. Thus, we performed parsimony ancestral state reconstruction for the parsimony data set, using the Most Parsimonious Reconstruction (MPR) algorithm in the R package ape for this data set. We then limited the data set to the 3,112 species nesting in only one type among hole, cup, or dome nests (hereafter the “likelihood” data set). We estimated transition rates among the three nest types by maximum likelihood under four different rate models (Pagel 1994; Paradis *et al*. 2004), and used AIC values to compare models. We estimated ancestral states using the make.simmap function in the R package phytools (Revell 2012) under the “all-rates-different” (ARD) model, which was preferred by AIC.

### Species ranges

We downloaded the BirdLife International/NatureServe (NatureServe 2014) range maps for passerine species on September 18, 2015, and examined latitudinal variation among species ranges using R 3.3.3 {{331 Core 2012;}}. To prevent errors from invalid geometries in species ranges, we first cleaned breeding range polygons by polygonation using the function clgeo_Clean (package cleangeo 0.2-2, https://github.com/eblondel/cleangeo). We then calculated the centroid of the breeding range using the function gCentroid (package rgeos 0.3-23 http://r-forge.rproject.org/projects/rgeos/).

For elevation analyses, we limited our species data set to the 874 passerine species in our “PGLM” data set (see below) whose range centroids were within 23.433°S and 23.433°N and –30° and -130°W, and whose elevational range could be estimated with our data set. This data set is hereafter referred to as the “Neotropical passerine” data set. We calculated the median elevation for each species’ breeding range by first subsetting a digital elevation model (DEM) by the shape of the breeding range, resulting in a DEM with the same limits as the breeding range. We then calculate the median elevation of the pixels across the entirety of the breeding range, using the cellStats function in the R package *raster* 2.5-8. To obtain a DEM covering all of the western hemisphere, we combined country-level DEMs available through the *raster* function getData (also available at: http://www.diva-gis.org/gdata). These DEMs are aggregated at a resolution of 30 seconds from a CGIAR SRTM 3-second resolution DEM (Reuter *et al*. 2007).

### Body mass data

We associated body mass data from a large compendium of avian body masses (Dunning 2008, 2015) with the species that were in both our nesting data set and phylogeny. Where separate body mass estimates are made for males and females in this data set, we took the average of the male and female body mass. Taxonomic reconciliation was required to match some mass data with tips in the Burleigh et al. (2015) phylogeny, and to the nest data from the Handbook of the Birds of the World Alive and species range data. We reconciled names by checking for species names from the body mass data set without matches in the other data, and examining taxonomic history to check for synonyms as above (see “Phylogeny” section). We matched body mass data and species range data with 2,754 of the 3,112 species in the “likelihood” data set, yielding a new data set which we refer to as the “PGLM” (phylogenetic generalized linear model) data set.

### Phylogenetic generalized linear models

To analyze latitudinal and elevational variation in the probability of evolving different nest types, we used phylogenetic generalized linear models. We built simple models akin to logistic regression, and accounted for phylogenetic effects by modeling the evolution of nest type with a threshold model where an underlying continuous value evolves under Brownian motion. We ran different models for each nest type (hole, cup and dome) using the R package phylolm (Ho & Ane 2014). The response variable in each of these models was the nest type of interest versus all other nest types (e.g. dome-nesting versus not dome-nesting). We could not perform phylogenetic logistic regression with three response categories, representing each nest type, as it has not been implemented in the framework we used for analysis (Ho & Ane 2014). For latitudinal analyses, our full model included the absolute value of latitude, log body mass, and their interaction as predictors. For elevational analyses on the Neotropical passerine data set (minus 14 species that did not have body masses reported: Dunning, 2008; Dunning, 2015), we built models with median elevation, log body mass, and their interaction as predictors.

We also investigated whether nest type and latitude (or elevation) predicts log body mass, instead of predicting nest type with log body mass. We performed this analysis because nest type is strongly conserved within lineages (Figure 1), and thus nest type may define evolutionary regimes for the evolution of log body mass, instead of responding to body mass and latitude (or elevation). For these analyses, we built phylogenetic generalized linear models with log body mass as the response variable, and with nest type and latitude (or elevation) and their interaction as predictors. These models allow us to contrast relationships of log body mass across ecological gradients within each nest type. We again implemented these models using phylolm. These models are built under the assumption that the evolution of log body mass is adequately described by an Ornstein-Uhlenbeck process (using the OUrandomRoot option in model calls).

**Figure 1 legend.**
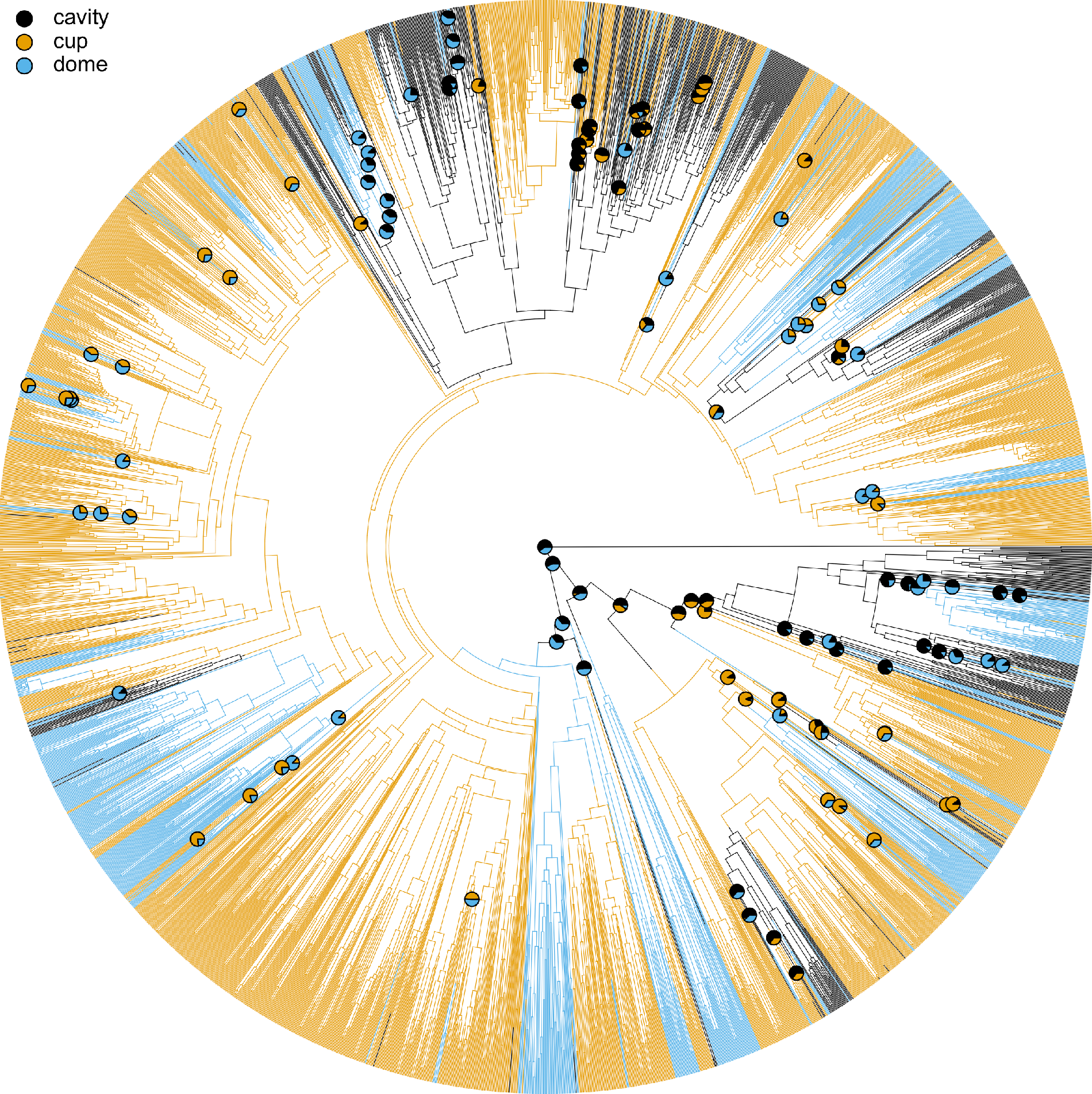
A stochastic reconstruction of nesting behavior across the passerine phylogeny under the all-rates-different (ARD) model detailed in Table 2, including 3,122 species as tips in the phylogeny (a subtree of the Burleigh et al. 2015 maximum likelihood phylogeny). Note that transitions are relatively rare, with many large clades dominated by a single nest type. Pie charts at nodes indicate posterior probabilities for each nesting types at all nodes where the maximum posterior probability for any nest type was < 0.9. Note that the ancestral nesting state for birds is reconstructed as either domed or cavity (see Price and Griffiths 2017).

## Results

### Nest type prevalence

In the “all species” data set comprising the 4,373 passerine species that could be scored for nest type or nesting behavior, 27 (0.62%) are brood parasites that do not construct a nest or incubate eggs. Of the species that construct a nest or incubate eggs (4,346 species), 560 (12.9%) are hole nesters, 2,546 (58.6%) are open cup nesters, and 1,117 (25.7%) are dome nesters. The remaining 123 species are scored as nesting in more than one nest type: hole or cup, 65 species (1.5%); hole or dome, 17 species (0.39%); cup or dome, 41 species (0.94%).

### Ancestral state reconstructions

Both maximum parsimony and maximum likelihood ancestral state reconstructions of nesting behavior show that the cup, dome, and hole nesting states are strongly conserved across most of the passerine phylogeny. Transitions among nest types are rare in both the maximum parsimony (Supplementary Information Figure 1) and maximum likelihood (Figure 1) reconstructions. Many large clades are dominated by a single nest type. Among the transition rate models in maximum likelihood reconstructions, the preferred model by AIC is the ARD (all-rates-different) model (Table 1). The transition rates in this model are low (see Table 2; all transition rate categories ≤ 0.0104 per million years while tree height = 56.9 million years).

**Table 1:** AIC scores for evolutionary transition models in nesting type and gregariousness (see Methods).

| | AIC | ΔAIC |
| --- | --- | --- |
| ***Nesting behavior*** |  |  |
| **All rates different** | 2120.65 | 0 |
| **Equal rates** | 2203.72 | 83.07 |
| **Symmetric** | 2204.08 | 83.43 |
| **Symmetric hidden and open** | 2205.45 | 84.80 |

**Table 2:** Evolutionary transition rates between nesting states from the ARD (all-rates-different) model for nesting behavior states. Overall transition rates are low (see Fig. 1). Estimated transition rates to open cup nesting from either cavity or dome nesting were greater than the reverse.

| Transition | Estimated rate (± SE) |
| --- | --- |
| **cup -> hole** | 0.001238 ± .0001945 |
| **dome -> hole** | 0.00186 ± .0004542 |
| **hole -> cup** | 0.01039 ± .001095 |
| **dome -> cup** | 0.004109 ± .0005852 |
| **hole -> dome** | 0.006875 ± .0009814 |
| **cup -> dome** | 0.003192 ± .0002882 |

Despite the relatively low frequency of dome and hole nesting among extant taxa, ancestral state reconstruction by maximum likelihood consistently finds that the most recent common ancestor (MRCA) of all extant passerine lineages nested in either domes or cavities (Figure 1). Our maximum likelihood results are consistent with those of (Price & Griffith 2017), in that we recovered a nest type other than open cup as the state of the MRCA of the extant passerines. Meanwhile, the nest type state of the MRCA of the extant passerines in ancestral state reconstruction by maximum parsimony is more ambiguous (Supporting Figure 1), with all nest types possible in the two most basal nodes.

Our results suggest that the rarity of ‘roofed’ (hole and dome) nests as compared to open cup nests can be explained in part by transition rate biases that favor transitions to open cup nesting. Our estimated transition rates in maximum likelihood analyses are highly asymmetric between hole- and cup-nests, with the hole to cup rate nearly an order of magnitude higher than the cup to hole rate (0.010 transitions versus 0.0012 transitions per million years, respectively, Table 2). Similarly, transitions to open cup nesting from dome nesting were estimated to occur at a ~25% higher rate than the reverse (Table 2).

### Nest type by latitude

Under the hypothesis that roofed nests gain thermoregulatory benefits through slower heat dissipation from nest contents (eggs, nestlings, and/or the incubating adult), we predicted that roofed nests should disproportionately be found at high latitudes, where nest contents are more likely to be subjected to colder weather. However, the latitudinal pattern of nest use among species runs counter to this prediction (Figure 2 and Supplementary Figure 2). Dome-nesting species are predominantly found at low latitudes (Figure 2; see also Collias and Collias 1984, Auer *et al*. 2007, Martin et al. 2017). Assuming that predation rates are higher at low latitudes (Skutch 1985), the prevalence of dome-nesting species at low latitudes is more consistent with predictions from predation rate variation across latitudes (Martin 1995). There is a steep decline in species diversity of dome-nesting passerines at ~35° latitude compared to the diversity at lower latitudes (Figure 2, Supplementary Figure 2). In contrast to dome-nesting species, the relative prevalence of hole-nesting across species appears to have a subtle mid-latitude peak (Figure 2). In the northern hemisphere especially, moderate levels of hole-nesting species diversity are maintained past 40° N (Supplementary Figure 2). The proportion of species range centroids belonging to hole-nesting species appears highest near 40° N, although these proportions are fairly flat across latitude (Figure 2).

**Figure 2 legend.**
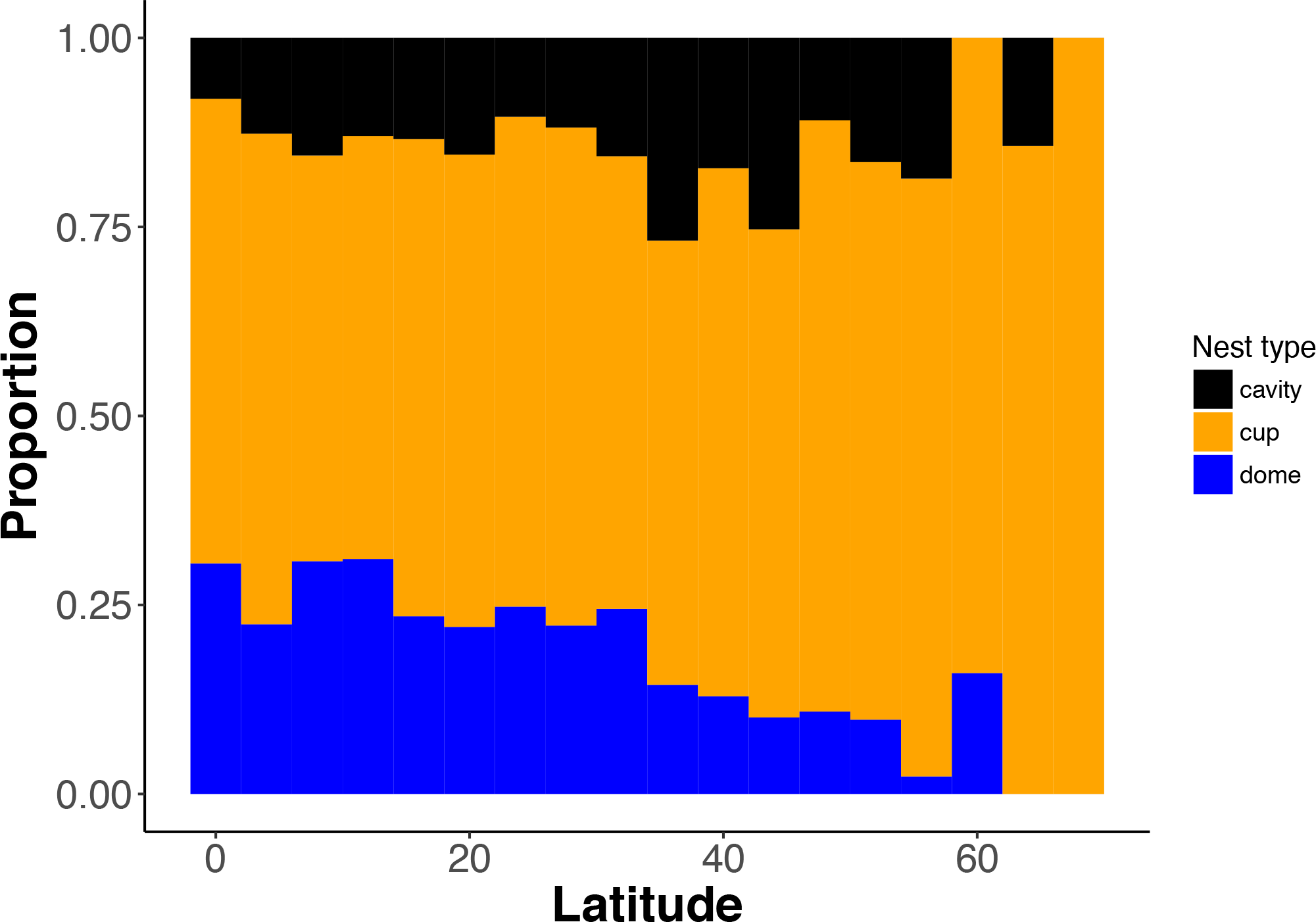
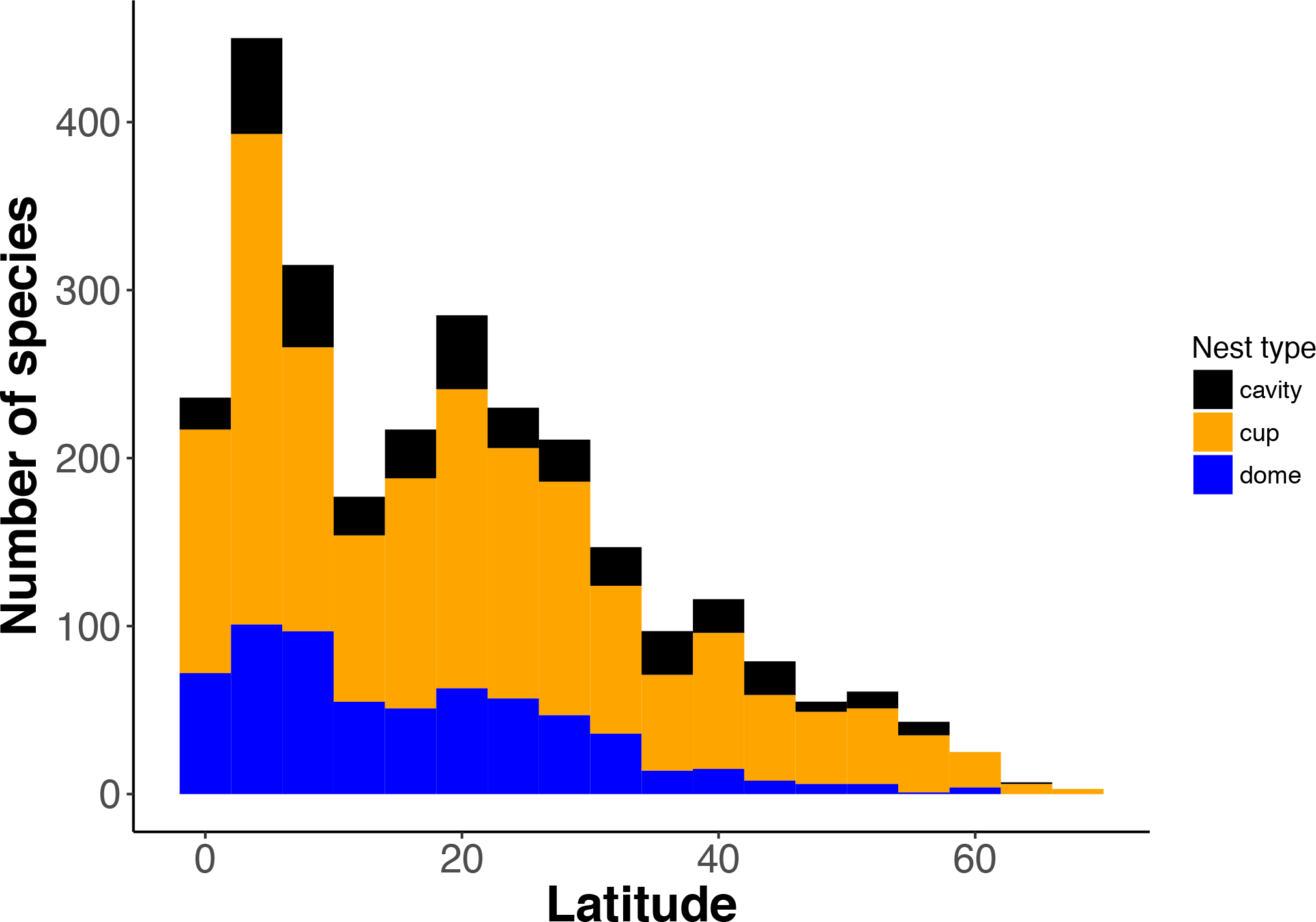
**a**. Proportions of species range centroids by latitude among 2,754 passerine species. The centroids falling within each latitudinal interval are used to calculate these proportions. **b.** Histogram showing the number of species range centroids for each nest type within latitudinal intervals.

In phylogenetic generalized linear models built to investigate whether latitude and body mass influence the evolution of cup-, dome-, and hole-nesting, intercept-only models were preferred for all three nest types by AIC (Table 3). This result indicates that neither latitude nor body mass is predictive of the evolution of the three different passerine nest type categories used in this study.

**Table 3:** Model selection for generalized linear models analyzing the probability of evolving different nest types across the passerines (n = 2,754 species), with latitude and log body mass as predictors.

| Response variable | Predictors | AIC |
| --- | --- | --- |
| Prob (cup-nesting) | centroid latitude, log body mass, interaction | 1497.147 |
| Prob (cup-nesting) | centroid latitude, log body mass | 1416.523 |
| Prob (cup-nesting) | centroid latitude | 1420.480 |
| Prob (cup-nesting) | log body mass | 1370.852 |
| Prob (cup-nesting) | none (intercept only) | **1368.844** |
| Prob (dome-nesting) | centroid latitude, log body mass, interaction | 1379.635 |
| Prob (dome-nesting) | centroid latitude, log body mass | 1478.349 |
| Prob (dome-nesting) | centroid latitude | 1231.565 |
| Prob (dome-nesting) | log body mass | 1193.085 |
| Prob (dome-nesting) | none (intercept only) | **1191.040** |
| Prob (cavity-nesting) | centroid latitude, log body mass, interaction | 872.7737 |
| Prob (cavity-nesting) | centroid latitude, log body mass | 857.2534 |
| Prob (cavity-nesting) | centroid latitude | 865.0302 |
| Prob (cavity-nesting) | log body mass | 855.4775 |
| Prob (cavity-nesting) | none (intercept only) | **853.1694** |

### Nest type by elevation

In the Neotropical species data set, there are no clear patterns of nest type prevalence with elevation. However, the great majority of Neotropical species ranges have low median elevations (<1000 m), such that the species diversity of all Neotropical passerine species is low at high elevation. High-elevation species diversity is especially low in the dome- and hole-nesting species (Figure 3a). Thus, range analyses did not provide evidence that either dome or hole nests are disproportionately prevalent at higher elevations in the Neotropics.

**Figure 3 legend.**
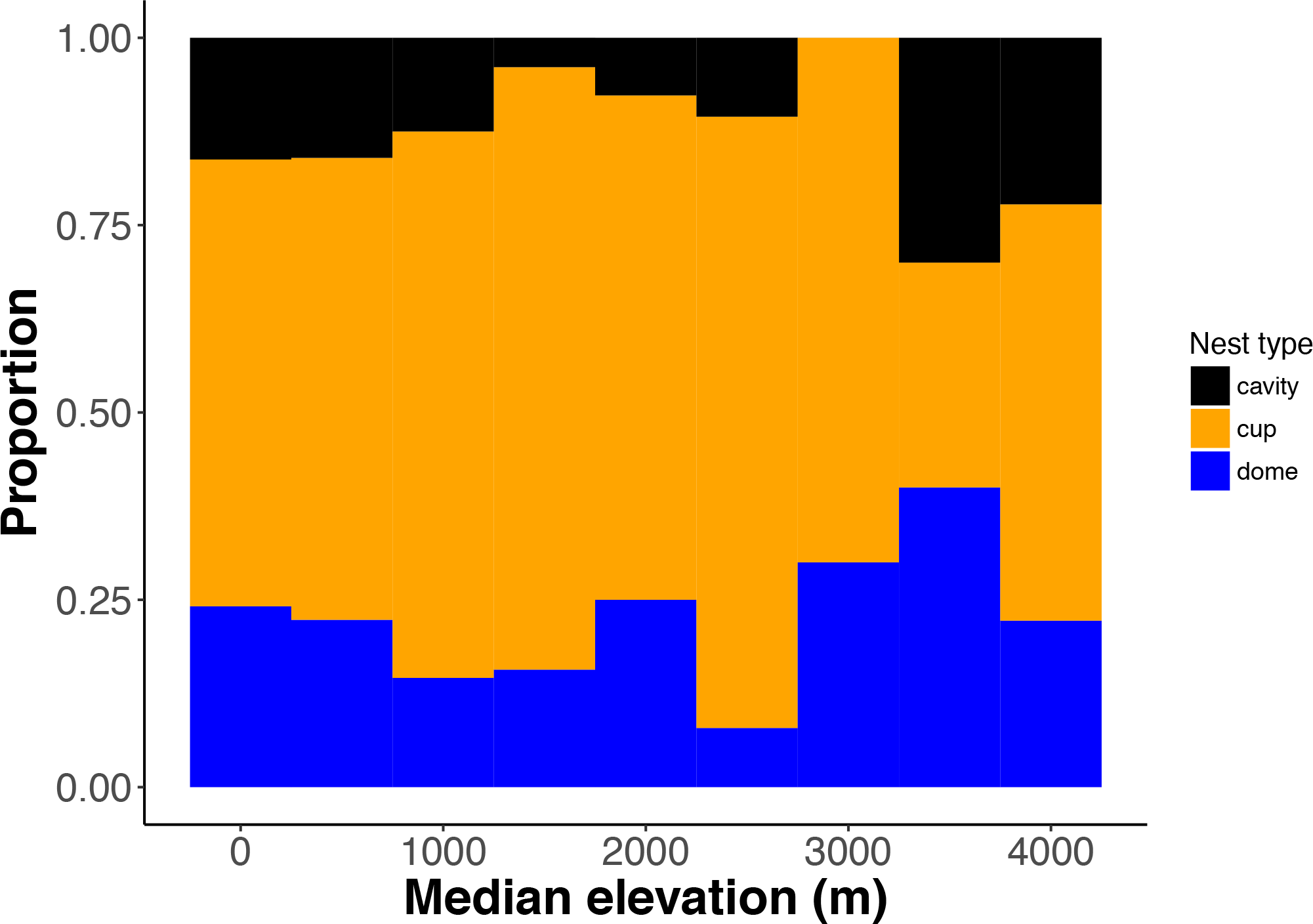
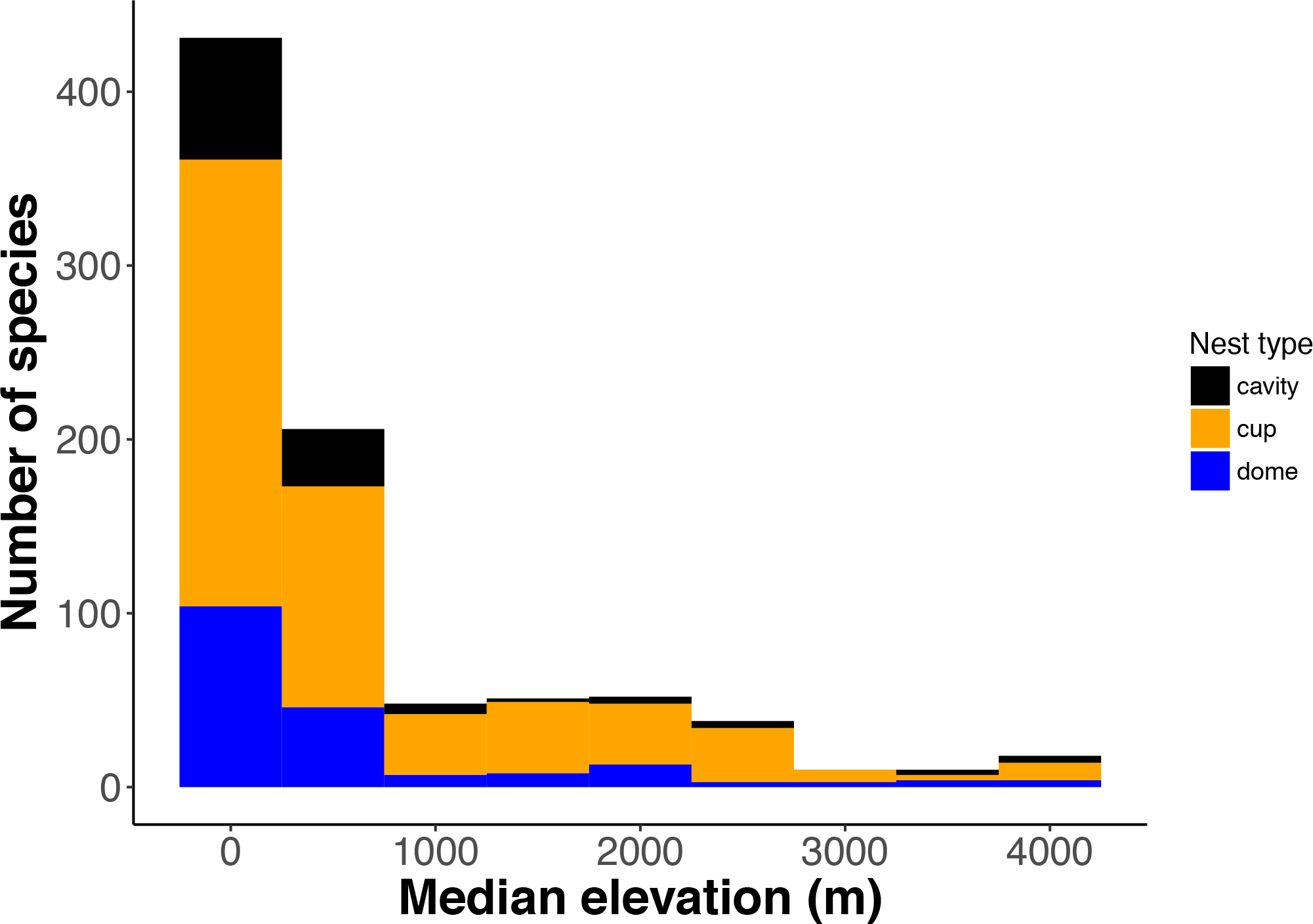
**a**. Proportions of species range centroids by elevation among 874 Neotropical passerine species. The centroids falling within each interval are used to calculate these proportions. **b**. Histogram showing the number of species range centroids for each nest type within intervals.

In candidate sets of phylogenetic generalized linear models built to test for the combined effects of elevation and body mass on the evolution of nest type, the preferred models by AIC for open cup-nesting and dome-nesting included only elevation as a predictor (Table 4). In the cup-nesting model, the probability of evolving cup-nesting increases slightly with median elevation (Table 5). However, the confidence intervals around the elevation parameter estimate, obtained from bootstrapping (Ho & Ane 2014), include zero. In the dome-nesting model, the probability of evolving dome-nesting decreases slightly with median elevation. The confidence intervals around the median elevation parameter again include zero in the dome-nesting model. The point estimates for the effect of median elevation in these models are consistent with expectations from the predation hypothesis, and inconsistent with expectations from the thermoregulation hypothesis. However, due to the uncertainty around the parameter estimates, these results should not be viewed as especially strong evidence for the correlations that should arise under the hypothesis that nest predation rates influence geographic ranges of the different nest types. For hole-nesting, the intercept-only model was preferred by AIC (Table 4).

**Table 4:** Model selection for generalized linear models analyzing the probability of evolving different nest types for neotropical passerines (n = 846 species).

| Response variable | Predictors | AIC |
| --- | --- | --- |
| Prob (cup-nesting) | Log mean elevation, log body mass, interaction | 499.2075 |
| Prob (cup-nesting) | Log mean elevation, log body mass | 502.0388 |
| Prob (cup-nesting) | Log mean elevation | **496.1432** |
| Prob (cup-nesting) | log body mass | 497.729 |
| Prob (cup-nesting) | none (intercept only) | 496.2275 |
| Prob (dome-nesting) | Log mean elevation, log body mass, interaction | 472.7101 |
| Prob (dome-nesting) | Log mean elevation, log body mass | 468.0592 |
| Prob (dome-nesting) | Log mean elevation | **466.2324** |
| Prob (dome-nesting) | log body mass | 467.7962 |
| Prob (dome-nesting) | none (intercept only) | 467.2081 |
| Prob (hole-nesting) | Log mean elevation, log body mass, interaction | 293.4847 |
| Prob (hole-nesting) | Log mean elevation, log body mass | 285.4204 |
| Prob (hole-nesting) | Log mean elevation | 283.4731 |
| Prob (hole-nesting) | log body mass | 283.4706 |
| Prob (hole-nesting) | none (intercept only) | **281.1440** |

**Table 5:**
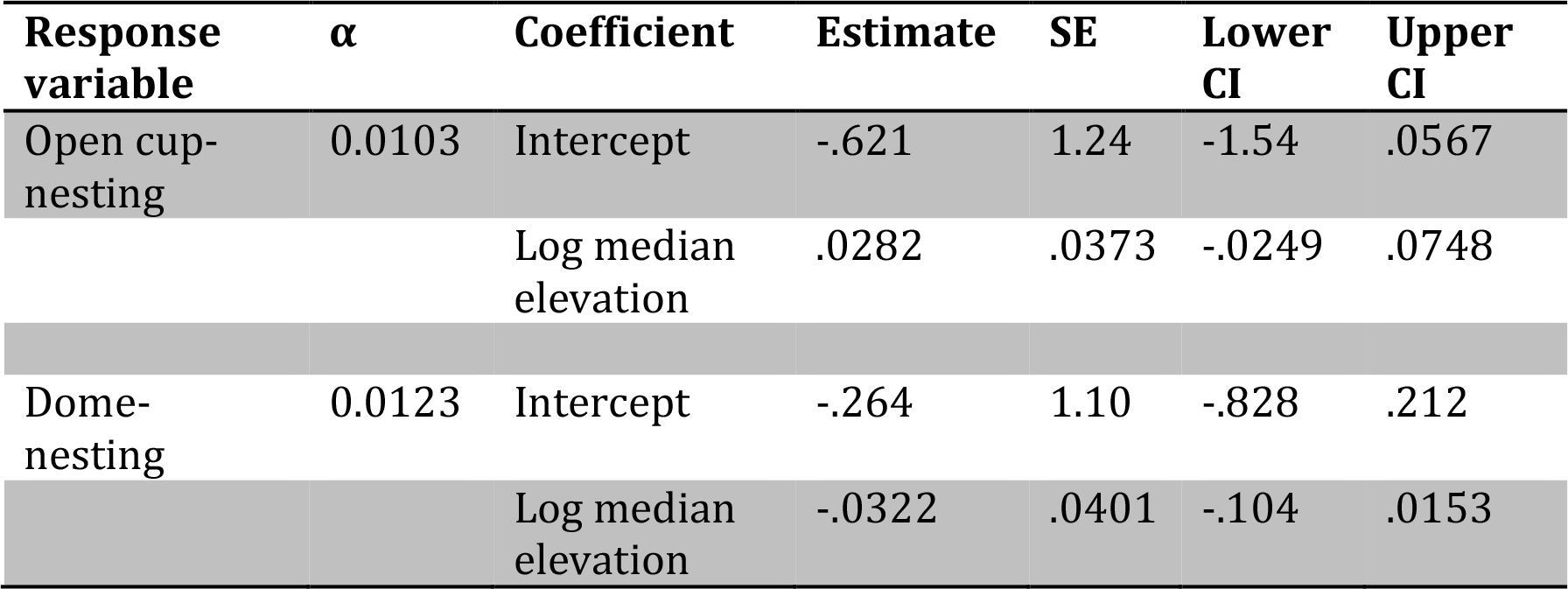
Coefficient estimates and uncertainty for preferred phylogenetic generalized linear models of the probability of evolutionary transition to open cup- and dome-nesting respectively. α is the rate of change in the continuous trait in the Brownian motion threshold model used in the model. It was not possible to estimate confidence intervals on the intercept (estimate = -.7439, SE = 1.4840, alpha = 7.80 x 10^−3^) of the cavity-nesting model.

### Predicting body mass by nest type and ecological gradients

In phylogenetic generalized linear models examining the evolution of body mass by nest type and latitude, the full model was preferred by AIC (Table 6). Visualization of the model results shows a complex pattern. Overall, dome-nesting species are smaller than both open cup- and hole-nesting species, consistent with previous evidence (Collias 1997; Martin *et al*. 2017). The most pronounced body mass differences by nest type are those between hole- and dome-nesting species in the tropics: hole-nesting species are approximately 2.3 times as large as dome-nesting species at the equator on average (based on expected values from phylogenetic GLM). While the body masses of cup- and dome-nesting species vary little with latitude, the body masses of hole-nesting species decline with latitude. The negative relationship of log body mass with latitude runs counter to the among-species interpretation of Bergmann’s rule (Olson *et al*. 2009), under which we expect a positive relationship.

**Table 6:** Model selection for generalized linear models analyzing the evolution of log body mass across the passerines (n = 2,848 species), with nest type and absolute latitude as predictors.

| Response variable | Predictors | AIC |
| --- | --- | --- |
| Log body mass | nest type, absolute latitude, interaction | **6636.691** |
| Log body mass | nest type, absolute latitude | 6660.258 |
| Log body mass | nest type | 6667.255 |
| Log body mass | absolute latitude | 6864.154 |
| Log body mass | none (intercept only) | 6863.386 |
| Log body mass | nest type, log mean elevation, | 888.6484 |
| Log body mass | nest type, log mean elevation | 885.0993 |
| Log body mass | nest type | 883.2454 |
| Log body mass | log mean elevation | 883.0425 |
| Log body mass | none (intercept only) | **881.1852** |

In phylogenetic generalized linear models examining the evolution of body mass by nest type and median elevation in neotropical passerines, the preferred model was the intercept-only model (Table 6). Thus, despite the negative relationship between log body mass and latitude found among cavity-nesting species, our study provides no evidence for a similar negative relationship between log body mass and elevation among hole-nesting species.

## Discussion

### Macroevolutionary dynamics of passerine nest types

Our combined macroevolutionary and macroecological analyses underscore the importance of evolutionary history in explaining the distribution of behavioral traits along ecological gradients. Associations of nest types with particular environments could arise from local adaptation if nest types can readily evolve to different environments. Alternatively, these associations could arise as an epiphenomenon, where clades dominated by particular nest types diversify at different rates at different places along latitudinal or elevational gradients. Our ancestral state reconstructions indicate that nest type states are generally conserved across the passerine phylogeny. Thus, the macroecological patterns of nest types are more likely a product of these macroevolutionary dynamics than resulting from local adaptation along gradients.

Although estimated transition rates between nest types were low (Table 4), the highest estimated rates of change were in hole-nesting lineages. This rate (< one per 50 million years per lineage) is low relative to what would be expected from regular nest type transitions via local adaptation, but it is noteworthy that hole-nesting species have the highest rates of transition among the nest types, as previous authors have hypothesized that hole nesting might constrain lineages from transitioning to other nest types (Collias & Collias 1984). The lowest rates of change (< one per 200 million years per-lineage) were in open cup-nesting species. Corresponding with these low transition rates, the evolutionary origin of nest type for most passerine species is ancient: ~97% of species in our data set trace the origin of their nest type back further than 10 million years (Figures 1 and 4). Open-cup nesting species in particular appear to have ancient origins for their nest types (Figure 4). While there are a few clades comprising tens of species that exhibit multiple transitions (e.g. the families Icteridae (Drury & Burroughs 2015), Muscicapidae (Barve & Mason 2015), Furnariidae (Zyskowski & Prum 1999; Irestedt *et al*. 2006), Timaliidae (Hall *et al*. 2015)), several large clades exhibit uniformity, or near-uniformity, in nest type.

**Figure 4 legend.**
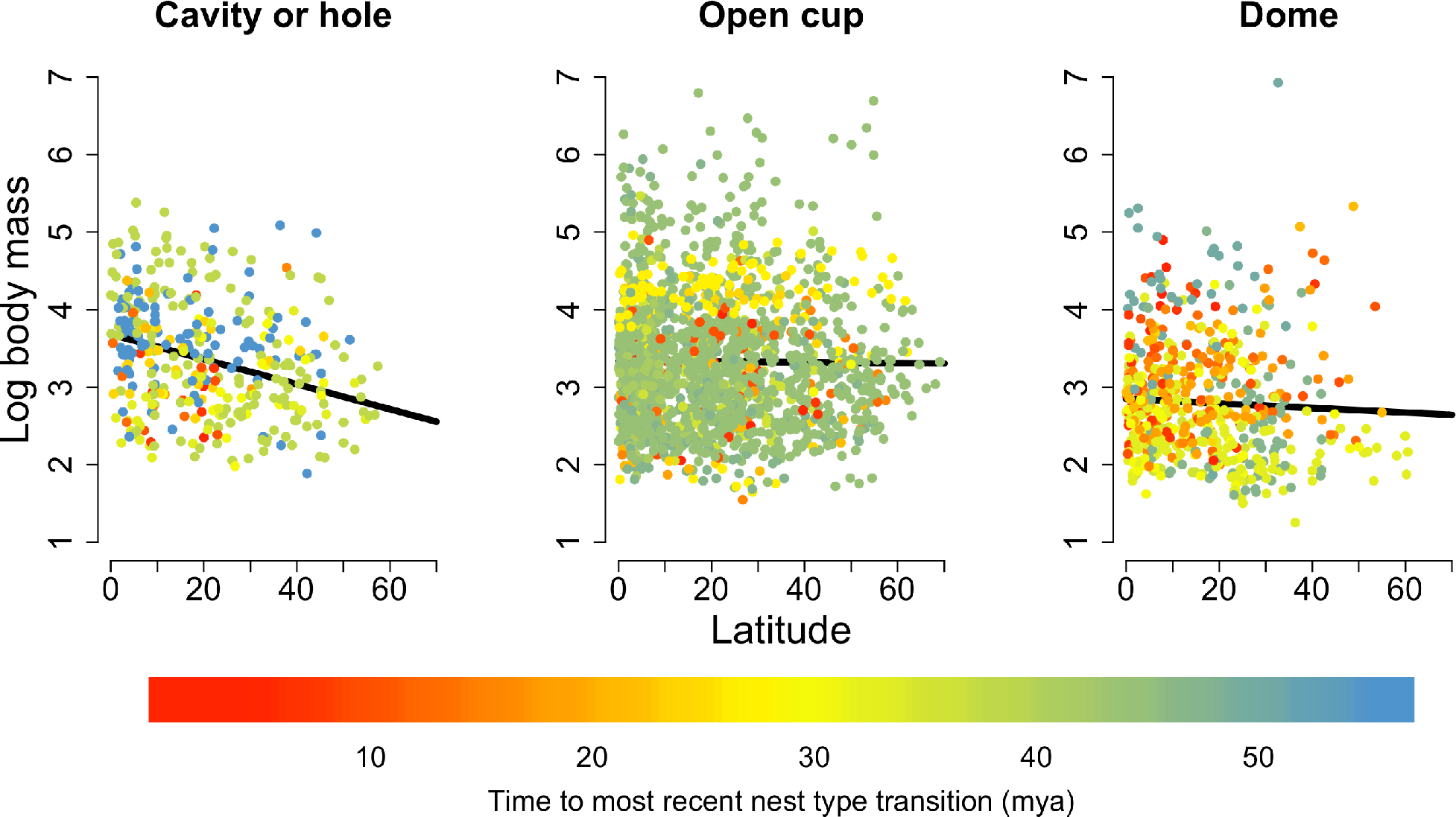
Patterns of log body mass across latitude by nest type for 2,754 species of passerine birds. Lines are predictions from phylogenetic generalized linear models, and are made irrespective of phylogenetic position of the data. Points are colored by time to most recent nest type transition) as estimated from a maximum likelihood ancestral state reconstruction for the PGLM data set. To estimate these times, we found the most recent node with a posterior probability <0.5 for a nest type different than the tip state for the species.

Some previous studies (Snow 1978; Price & Griffith 2017) have interpreted the evidence from passerines as exemplifying evolutionary conservatism in nest type as we do; however, Hansell (2000) interpreted previous evidence that multiple nest types exist within several passerine families as supporting evolutionary lability (see also Ligon, 1993), in arguing against hard constraints on nest type evolution. The number of transitions across the passerine phylogeny and the higher rates of change in some clades indeed indicate that there are no hard constraints on nest type evolution, but the substantial conservatism of passerine nest types (Table 2, Figure 4) stands in contrast to long-standing assertions that behavioral traits are generally prone to rapid evolutionary change (Darwin 1874; Blomberg *et al*. 2003). Further, such conservatism provides evidence that the time that most lineages have spent using any of the three nest types is adequate to enable extensive adaptive refinement of related life history traits (Lack 1947, 1954; Snow 1978; Martin & Li 1992; Bosque & Bosque 1995; Martin 1995; Auer *et al*. 2007; Barve & Mason 2015), egg color (Oniki 1985; Weidinger 2001), or other phenotypes under selection pressures that consistently differ by nest type.

Our ancestral state reconstructions indicate that all three nest types were represented in early passerine lineages (Figure 1), in accord with Collias (1997) and Price and Griffiths (2017). Because all three nest types were present in the early passerine lineages, Collias (1997) was skeptical that the nest type of the most recent common ancestor of the passerines could be known. Indeed, in both maximum likelihood and maximum parsimony reconstructions, the nest type state of the MRCA of the passerines represented in our phylogenetic tree is ambiguous. It is reconstructed as either cavity or dome-nesting in the maximum likelihood reconstruction (Figure 1), with open cup nesting secondarily evolved. In the maximum parsimony reconstruction, we achieve no resolution in the reconstruction of the MRCA among the three types (Supplementary Figure 1). However, even had we found an unambiguous reconstruction for the MRCA of crown Passeriformes in this analysis, skepticism is warranted for such reconstructions, as the extinction of even a single species in the early history of a clade can greatly affect the reconstructed states. That said, our maximum likelihood reconstructions agree with those of Price and Griffiths (2017) that open cup nesting is a derived state with respect to the crown Passeriformes, despite being the most common passerine nest type. Together with the macroecological patterns, which highlight the degree to which open cup nests are utilized across latitudinal and elevational gradients, these results raise the issue of why the open cup nest has become the most prevalent nest type. Further, they raise the question of how open-cup nesting clades have colonized a broader spectrum of environments than hole- and dome-nesting clades, including high-latitude and high-elevation areas, despite comparatively less protection from heat loss, exposure to sun and rain, and predation.

### Nest type prevalence along latitudinal and elevational gradients

If roofed nests have thermoregulatory benefits via reduced heat loss rates in cold environments, we should observe a greater prevalence of these nests at high latitudes, and at high elevation in the tropics. Our macroevolutionary evidence suggests that such a pattern, if it existed, would result from greater success of roofed-nesting lineages in the colonization of cold environments. However, the combination of the latitudinal and elevational patterns of species ranges do not provide substantial evidence that roofed nests aid lineages in the colonization of cold environments. The patterns are more clear for dome-nesting species than hole-nesting species. We find, in agreement with previous authors (Collias & Collias 1984; Martin *et al*. 2017), that dome-nesting species are more limited to the tropics than either hole-nesting or cup-nesting species, and that dome-nesting species make up a greater proportion of species diversity in the subtropical and temperate southern hemisphere than the northern hemisphere (e.g. Supplementary Figure 2). Our analyses add an important additional insight regarding the ranges of dome-nesting species: they are no more prevalent at high elevation than at low elevations within the Neotropics. Indeed, our analyses of the evolution of dome-nesting provide limited support for a negative relationship between dome-nesting and elevation – that is, our analyses show that evolutionary transitions to dome-nesting may be more likely at lower elevations. Whatever thermoregulatory benefits come from using a domed nest, these benefits have not resulted in transitions to dome nests in cold environments, or disproportionately predisposed lineages using domed nests to successful colonization of and diversification within colder environments. Our analyses instead reinforce the degree to which dome-nesting species are concentrated in the lowland tropics (Figure 3b). Thermoregulatory advantages may be important to other aspects of dome-nesting passerine biology, such as permitting longer durations of off-nest activities during incubation because of lower rates of cooling (Martin et al., 2017). The importance of such an advantage would seem to be lowest, however, in the lowland tropics, where cooling of eggs and nestlings is slow because of ambient temperatures. It is also possible that the roof’s benefits may come from shielding nest contents from sun or rain exposure (Snow 1978), two likely challenges for nesting birds in the lowland tropics, and especially for birds that spend long periods of time away from the nest (White & Kinney 1974; Deeming & Gray 2016).

The distribution of dome-nesting species along latitudinal and elevational gradients is consistent with the hypothesis that higher predation risk in the lowland tropics renders dome-nesting a more successful strategy there than elsewhere. This interpretation assumes that predation risk has consistently been higher at lower latitudes (Skutch 1966, 1985; Snow 1978) and lower elevations (Boyle, 2008; Jankowski et al., 2013; Skutch, 1985) over evolutionary time, that the domed-nesting habit results in reduced nest predation rates compared to the open cup-nesting habit (Hall *et al*. 2015; Martin *et al*. 2017), and that a tradeoff exists that renders dome-nesting less advantageous at higher latitudes, despite its advantages with respect to predation. The first two of these assumptions appear plausible based on the results of existing studies (but see Martin et al. 2017), while the third has not, to our knowledge, been studied. With respect to the predation benefits of domed nests, Martin et al. (2017) focused on the inconsistency in the outcome of predation rate comparisons between open cup and domed nests. However, of ten within-site predation rate comparisons for open cup versus domed nests presented by Martin et al. (2017), nine yield higher predation rate estimates for open cup nests than domed nests (although not all nine of these within-site differences are statistically significant).

The distributions of hole-nesting species, and the relative prevalence of hole-nesting across latitude and elevation, are less clear. Compared with the dome-nesting species, hole-nesting species are less restricted to low latitudes, with their relative prevalence highest at mid-latitudes. Their relative prevalence does not correspond with the expectations of either the predation or thermoregulation hypotheses. Predation rates are lower in hole nests than in domed nests (Auer *et al*. 2007), so the relative benefits of protection from predation for hole-nesting should be strongest where predation rates are highest – at low latitude and low elevation – yet hole nests do not have their greatest prevalence there. Predation rates may also be higher in the southern hemisphere than the northern hemisphere (Martin 1996), but hole-nesting species diversity declines much more steeply with latitude in the southern than the northern hemisphere (Supplementary Figure 3), counter to predictions for the distribution of hole-nesting species from the predation hypothesis.

Local diversities of hole-nesting species may be limited by the availability of suitable nest sites, via limits on population densities (Newton 1998; Cockle *et al*. 2010). We might expect the greatest availability of hole-nesting sites in vast lowland forests like the Amazon and Congo basins, where there are so many trees in various stages of decay. Why do hole-nesting passerine species not have greater prevalence in the lowland tropics, then? One potential explanation is competition for cavities with non-excavating, non-passerine species like parrots and trogons (Brightsmith 2005) – clades that are largely confined to the tropics and sub-tropics. Thus, the mid-latitude peak in hole-nesting passerines may in part be explained by a competition gradient for nest sites from non-passerines. An alternative explanation is that tropical tree cavities decay or are colonized by parasites more quickly, such that the number of cavities in these forests is far greater than the number of suitable nest cavities (Lõhmus & Remm 2005; Cockle *et al*. 2010).

### Body size evolution in association with nest type

Our analyses of body mass evolution across latitude revealed an unexpected pattern: hole-nesting species become smaller at higher latitudes (Table 3, Fig. 4), but not at higher elevations. In our PGLM, the expected body mass for hole-nesting passerines is ~40 g at the equator, and ~18 g at 50° latitude. The relationship between body mass and latitude in hole-nesting species is counter to Bergmann’s rule *sensu lato*, which has some support across birds more generally (Olson *et al*. 2009). However, we do not recover any evidence for a similar decline in body mass with elevation in hole-nesting species. Meanwhile, our analyses predict that body mass in equatorial dome-nesting species is just 36% of the body mass in equatorial hole-nesting species. The difference between these values is higher in the tropics than outside the tropics. Indeed, despite dome-nesting species generally having smaller mass than either hole- or open cup- nesting species, consistent with previous studies (Collias & Collias, 1984; Martin et al., 2017), the predicted values for hole-nesting and dome-nesting species converge at high latitudes (Figure 4).

Life history aspects correlated with the hole-nesting habit may help explain these patterns. Hole nesting is associated with longer developmental periods (Martin & Li 1992). Further, developmental periods increase with body mass in passerines (Bosque & Bosque 1995). Thus, large-bodied hole-nesting species should generally have long developmental periods, with smaller-bodied hole-nesting species having shorter developmental periods. The length of the breeding season, meanwhile, decreases with increasing latitude (Ricklefs 1966; Barve & Mason 2015), but likely does not decrease with elevation to the same degree, if at all. It is therefore possible that large-bodied hole-nesting species cannot sustain population growth at high latitudes, resulting in a filtering of hole-nesting species by developmental time at higher latitudes, whereas no such filtering is evident at higher elevations. Short breeding seasons could limit the prospects for re-nesting following failure, or multiple clutches, which could limit population growth and hence colonization of high latitudes by larger hole-nesting species. Importantly, this argument relies on conservatism in life history traits associated with hole nesting, and a failure for local adaptation to drive faster development of hole-nesting species at higher latitudes. This issue requires further investigation. We note, however, that the short development times of open cup-nesting species might explain, in part, why open cup-nesting species dominate at the extreme high end of the latitude spectrum (Figure 2) where breeding seasons should be shortest.

## Acknowledgements

This work was funded in part by National Science Foundation grants DBI-1458034 and DEB-1208428, and by the University of Florida.

